# Altered environmental perception by parental stress and depression vulnerability: impact on mothers and offspring

**DOI:** 10.1101/2021.02.23.432580

**Authors:** Renata L. Alves, Camila C. Portugal, Igor M. Lopes, Pedro Oliveira, Cecília J. Alves, Fernando Barbosa, Teresa Summavielle, Ana Magalhães

**Author notes:** Corresponding author: Ana Magalhães.

## Abstract

Depressive mothers often find the mother-child interaction to be challenging. Parental stress may further impair mother-child attachment, which may increase the risk of negative developmental consequences. We used rats with different vulnerability to depression (Wistar and Kyoto) to investigate the impact of stress (maternal separation-MS) on maternal behaviour and adolescent offspring cognition. MS in Kyoto dams increased pup-contact, resulting in higher oxytocin levels and lower anxiety-like behaviour after weaning, while worsening their adolescent offspring cognitive behaviour. Whereas MS in Wistar dams elicited higher quality of pup-directed behaviour, increasing Brain-Derived Neurotrophic Factor (BDNF) in the offspring, which seems to have prevented a negative impact on cognition. Hypothalamic oxytocin seems to impact the salience of the social environment cues (as negative for Kyoto) leading to different coping strategies. Our findings highlight the importance of contextual and individual factors in the understanding of the oxytocin role in modulating maternal behaviour and stress regulatory processes.

## Introduction

Depressive mothers often display higher difficulty in interacting with their child, increasing the liability of child abuse and neglect ^1^. Given the increase in parental adverse events (e.g., pandemic, lockdowns, poverty, massive migration, displacement, etc), parental stress is augmented, which may translate to an increased rate of depressive mothers, and in turn, the rise the worldwide number of children growing up under chronic stress ^2^. Thus, parental stress may have long-lasting effects on mothers, their children, and the family. Despite the importance of vulnerability to depression and its relationship with parental stress (as an early stressful emotional event), there is a lack of research combining the impact of these two risk factors on mother’s behaviour and its consequences for the offspring.

In rodents, maternal behaviour allows a proper attachment between mother and pups, adequate nutrition, and pup’s temperature regulation ^3^. It includes a complex variety of behaviours and its quality and quantity have been connected to the development of emotional and cognitive processes in the offspring ^4^. Maternal behaviour is regulated by oxytocin ^5^, a neuropeptide released from the paraventricular nucleus and the supraoptic nucleus of the hypothalamus ^6^. Oxytocin activates brain areas related to maternal behaviour and its blockage leads to maternal care deficits ^7, 8^. Furthermore, the oxytocinergic system is impacted by motheŕs absence ^9^. The maternal separation paradigm (MS), used as a parental adverse event, is the most widely used animal model to study early life stress and disruptions of the infant-mother relationship ^10^. Under MS, pups increase solicitation calls ^11^, compelling mothers to increase their maternal behaviour as a response to pup needs ^4^, which has been associated with increased oxytocin receptors (OXTR) expression ^12^.

The scarce existing evidence seems to indicate that MS induces changes in the hippocampus of the offspring ^13, 14^, leading to maladaptive brain development, resulting in cognitive impairment in adulthood ^15^. Of note, psychosocial stressors can lead to increased microglial reactivity in the hippocampus, which may also contribute to cognitive impairment ^16^, as neuroinflammation is now accepted as a crucial mediator in cognitive impairment ^17^ Moreover, neuroinflammation is also linked to depression ^18^, primarily by its association with increased expression of proinflammatory cytokines, such as the tumour necrosis factor (TNF), interleukin (IL)-6 and IL-1β ^19–21^.

In the present study, we used a depressive-like animal model combined with the MS paradigm to investigate the interplay between the role of pre-existing vulnerability to depression and environment/social risk factors. Using the Wistar-Kyoto (Kyoto) strain, frequently used as a genetic model for endogenous depression-like susceptibility and the Wistar rat as a control strain ^22–24^, we investigated the impact of MS on maternal care, mother’s anxiety- and depressive-like behaviour with a specific focus on the interactions between the oxytocinergic system and neuroinflammatory processes. Evaluation of MS impact on offspring was assessed by analysing the adolescent offspring performance in learning, spatial and working memory behavioural tests combined with two glutamatergic synaptic proteins expression in the hippocampus as well as neuroinflammation levels. Our results showed that depressive-like mothers had higher levels of oxytocin as a stress response mechanism, which translated into the perception of the environment as insecure. As a result, they kept their pups closer to them and away from outsiders. The non-depressive-like mothers (Wistar) maternal behaviour (more affiliative) observed after MS seems to protect the cognitive performance of adolescent offspring, with an increase of hippocampus BDNF. On the other hand, depressive-like adolescent offspring showed alteration in the IL-1β levels and worst performance in the cognitive tests, suggesting lower resilience to MS effects.

## Results

### Evaluation of mothers’ behaviour in response to maternal separation (MS)

#### Wistar dams presented higher-quality maternal behaviour than Kyoto dams in response to MS

MS impact on mothers and their offspring occurs due to the separation period but more importantly, due to changes in their interaction afterwards. Thus, we started by investigating the MS effect on maternal behaviour. We observed that the total time spent on maternal behaviour was higher in both MS groups (*F*_(1,26)_ = 15.7, *p* ≤ 0.001) independently of the strain. In both strains, MS dams spent significantly more time in maternal behaviour than control dams of the same strain (Wistar: *p* < 0.01; Kyoto: *p* < 0.05) **(Fig. 1A)**. However, the increased quantity of the time spent on maternal behaviour is not sufficient for the value of the mothers’ response to MS. As such, we analysed also the quality of maternal behaviour by grouping them in different types of maternal behaviour: affiliative behaviours (licking/grooming and nursing) or non-affiliative contact (time dams spend in contact with offspring excepted the affiliative behaviour) and observed an effect of strain for different behaviours. Affiliative behaviour was higher in MS Wistar dams than control Wistar dams (*p* < 0.01), which did not happen in the Kyoto strain (strain x MS: *F*_(1,26)_ = 3.93, *p* = 0.058; MS: *F*_(1,26)_ = 4.08, *p* = 0.054) **(Fig. 1B)**. In contrast, when we analysed the time spent in non-affiliative contact with pups, MS Kyoto dams spent more time in non-affiliative contact than control Kyoto dams (*p* < 0.001) and MS Wistar dams (*p* < 0.001) **(Fig. 1C)**. The impact of MS on non-affiliative contact was dependent on the strain (strain x MS: *F*_(1,26)_ = 14.0, *p* ≤ 0.001; MS: *F*_(1,26)_ = 4.95, *p* < 0.05). In order to further assess maternal care, pup weight over time was also analysed. Only a strain effect was observed (*p* < 0.001), with no MS effect for either strain (litter effect was considered as random effect) **(Fig. 1D)**. Taken together, control Wistar and Kyoto dams spent similar time in maternal behaviour. However, under MS conditions, the strains increased differently the time spent in maternal behaviour, with higher affiliative behaviour for Wistar and higher non-affiliative contact for Kyoto.

**Figure 1.**
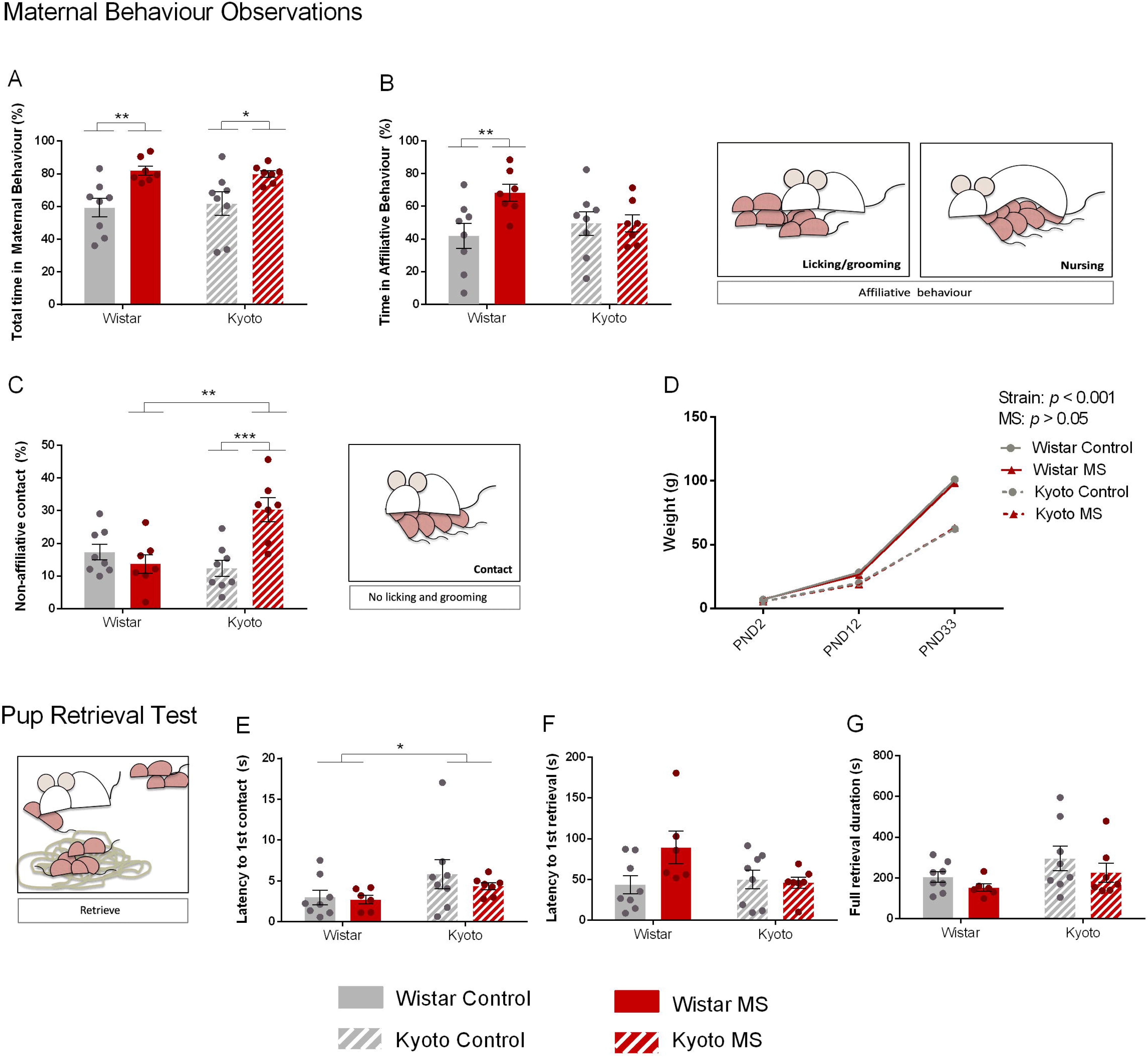
MS impacts maternal behaviour of mothers with different vulnerability to depression. (A) MS increased maternal behaviour in both strains. However, Wistar MS dams increased (B) affiliative behaviour (high-quality behaviours) and Kyoto MS dams increased (C) non-affiliative contact time. (D) Wistar showed higher weight than Kyoto strain, but weight was not impacted by MS. Data were analysed by two-way ANOVA (2 x 2 factorial design) with Bonferroni comparisons, except for (D) which was analysed with a linear mixed model (strain and MS as fixed effects and litter as random effect). (E) Latency to first contact was higher in the Kyoto strain. (A), (B) and (C) were data arcsine transformed. (E), (F) and (G) data were log-transformed. *n* = 8 (controls Wistar and Kyoto); *n* = 7 (MS Wistar and Kyoto, except for PRT: *n* = 6 for MS Wistar). Results are expressed as mean ±SEM. \**p* < 0.05, \*\**p* < 0.01, \*\*\**p* < 0.001. MS = Maternal Separation; PND = Postnatal day.

To further investigate maternal behaviour, we also performed the Pup Retrieval Test (PRT), revealing that Kyoto dams present a higher latency to the first mother/pup contact (strain: *F*_(1,25)_ = 4.55, *p* < 0.05) **(Fig. 1E)**. However, concerning the latency for the first pup retrieval and the total retrieval duration no differences were found **(Fig. 1F and 1G)**.

#### Increased exploration in Wistar dams during the MS period and decreased anxiety-like behaviour in Kyoto dams after the weaning period

Since an animal that experienced parental stress is expected to alter its emotional state, in addition to maternal behaviour, exploratory and anxiety-like behaviours by the mothers were also assessed. Anxiety-like behaviour in mothers in response to MS was evaluated using two different behaviour tests, the Elevated Plus Maze (EPM) and the Open-Field (OF) tests. The EPM was done during the MS protocol period while the OF test was conducted after weaning. Regarding the classical measurements for evaluating anxiety-like behaviour in the EPM, no strain or MS differences were found for the time spent or number of entries in the open arms **(Fig. 2A and 2B)**. However, we observed that Kyoto dams spent more time in the EPM centre area than Wistar dams, independently of MS (strain: *Wald χ*^2^ = 16.6, *p* < 0.001) **(Fig. 2C)** and that the Kyoto increased the total arm entries due to MS *(p* < 0.05) (MS: *F*_(1,26)_ = 7.55, *p* < 0.05) **(Fig. 2D)**. In contrast, Wistar dams displayed more rearing than Kyoto (strain: *F*_(1,26)_ = 50.4, *p ≤ 0.001, MS: F*_(1,26)_ = 4.22, *p* ≤ 0.05), with MS further increasing the Wistar number of rearing (p < 0.05) **(Fig. 2E)**. We observed the same pattern in relation to head dipping, with Wistar dams performing more head dipping than Kyoto dams (strain x MS: *Wald χ*^2^ = 7.32, *p* < 0.01; strain: *Wald χ*^2^ = 10.2, *p* ≤ 0.001; MS: *Wald χ*^2^ = 6.08, *p* < 0.05) and the MS impacting the frequency of head dipping only in Wistar (p < 0.001) **(Fig. 2F)**. Altogether, the results obtained in the EPM test suggest that anxiety-like behaviour was not affected during the MS protocol. However, Wistar dams displayed more exploratory behaviours than Kyoto dams and MS further increased exploration in Wistar.

**Figure 2.**
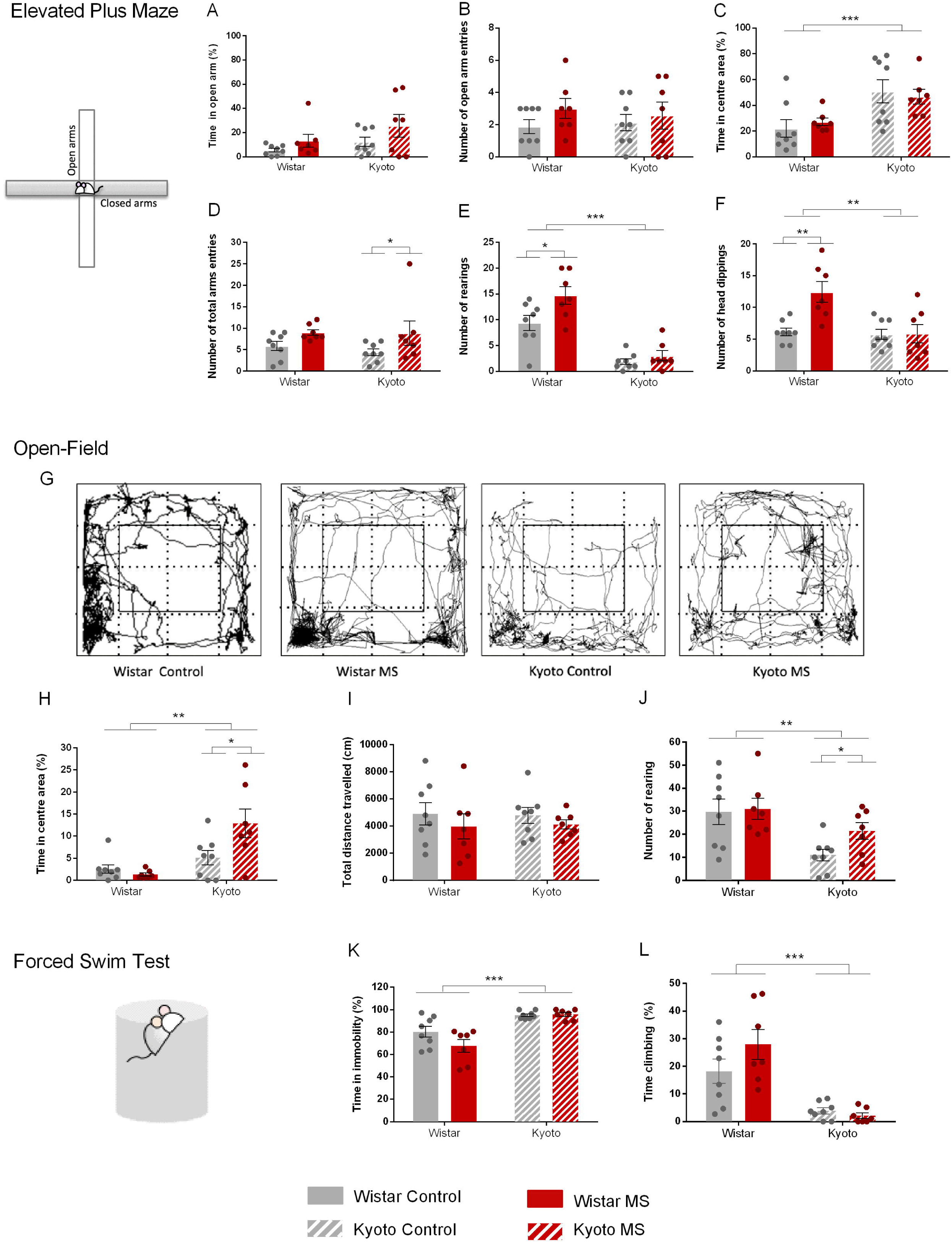
MS affects mothers’ emotional behaviour. During MS period (using EPM), (A), (B) no differences were found regarding anxiety. MS increased (D) Kyoto total arm entries and increased Wistar (E) number of rearing and (F) number of head dipping. After weaning (using OF), MS increased Kyoto (H) time in centre area and (J) number of rearing, indicating decreased anxiety levels. FST confirmed Kyoto depressive profile since Kyoto strain showed higher time (K) immobility and lower (L) climbing comparing with Wistar. MS did not impact depression-like behaviours. Data were analysed by two-way ANOVA (2 x 2 factorial design) with Bonferroni comparations. (A), (C), (H), (K), (L) data were arcsine transformed. (B), (D), (E), (F) data were log-transformed. (I), (J) data were square-root transformed. *n* = 8 (control and MS Wistar); *n* = 7 (control and MS Kyoto). Results are expressed as mean ±SEM. \**p* < 0.05, \*\**p* < 0.01, \*\*\**p* < 0.001. MS = Maternal Separation.

After weaning, exploratory and anxiety-like behaviours were studied with the OF test **(Fig. 2G)**. We observed that Kyoto dams spent more time in centre area than Wistar dams (strain x MS: *F*_(1,26)_ = 5.07, *p* < 0.05; *strain*: *F*_(1,26)_ = 12.1, *p* < 0.01). This was further reinforced by MS, withe MS Kyoto dams spending more time in the centre area than the control Kyoto (*p* < 0.05) and then MS Wistar dams (*p* ≤ 0.001). No MS effect was observed for Wistar **(Fig. 2H)**. However, total distance was similar in all groups **(Fig. 2I)**. Exploration behaviour, assessed as number of rearing, was different between the strains (strain: *F*_(1,26)_ = 11.4, *p* < 0.01). Globally, Kyoto dams presented fewer rearing in comparison to Wistar dams, yet MS Kyoto dams performed more rearing than control Kyoto dams (*p* < 0.05) **(Fig. 2J)**. The results obtained in the OF test suggest that Kyoto dams present less anxiety-like behaviour after weaning in comparison with Wistar dams and, of note, Kyoto dams subjected to MS displayed less anxiety-like behaviour than the control Kyoto dams. Exploratory behaviour results were similar during and after the MS protocol. Wistar dams displayed more exploratory behaviours than Kyoto dams, although MS increased exploration in Kyoto but not in Wistar dams.

#### MS did not influence depressive-like behaviour in dams

Considering our interest in depression and parental stress comorbidity, we investigated the effect of MS on both strains. Depressive-like behaviour in mothers in response to MS was evaluated using the Forced Swimming Test (FST). As expected due to their strain characteristics, Kyoto dams spent more time immobile in comparison with Wistar dams (*F*_(1,26)_ = 36.3, *p* < 0.001) **(Fig. 2K)**. Consistently, Kyoto dams also spent less time climbing than Wistar dams (*F*_(1,26)_ = 43.5, *p* < 0.001) **(Fig. 2L)**. These results indicated that Kyoto dams presented more depressive-like behaviours than Wistar dams, but surprisingly, MS did not influence the test results, which may be due to a ceiling effect since the Kyoto control already spent most of the test time immobile.

#### MS increased oxytocin expression in the hypothalamus of Kyoto dams

Since maternal behaviour is strongly influenced by oxytocin ^5, 8^, we analysed the expression of oxytocin in the hypothalamus of dams to clarify how exposure to MS would affect this neuropeptide ^6^. Surprisingly, we observed that Kyoto dams presented increased oxytocin levels in the hypothalamus compared with Wistar dams, and identified a strain-MS interaction effect (*F*_(1,16)_ = 6.60, *p* < 0.05). Interestingly, MS Led to increased oxytocin levels in Kyoto dams (*p* < 0.05) **(Fig. 3A)**. Because altered levels of oxytocin may result in altered expression of OXTR in oxytocin projection regions, we evaluated the expression of this receptor in oxytocin target regions that are recognised as relevant for regulation of maternal behaviour, such as the PFC, the hippocampus, and the amygdala. As represented in **Fig. 3B-3D,** no significant differences were observed.

**Figure 3.**
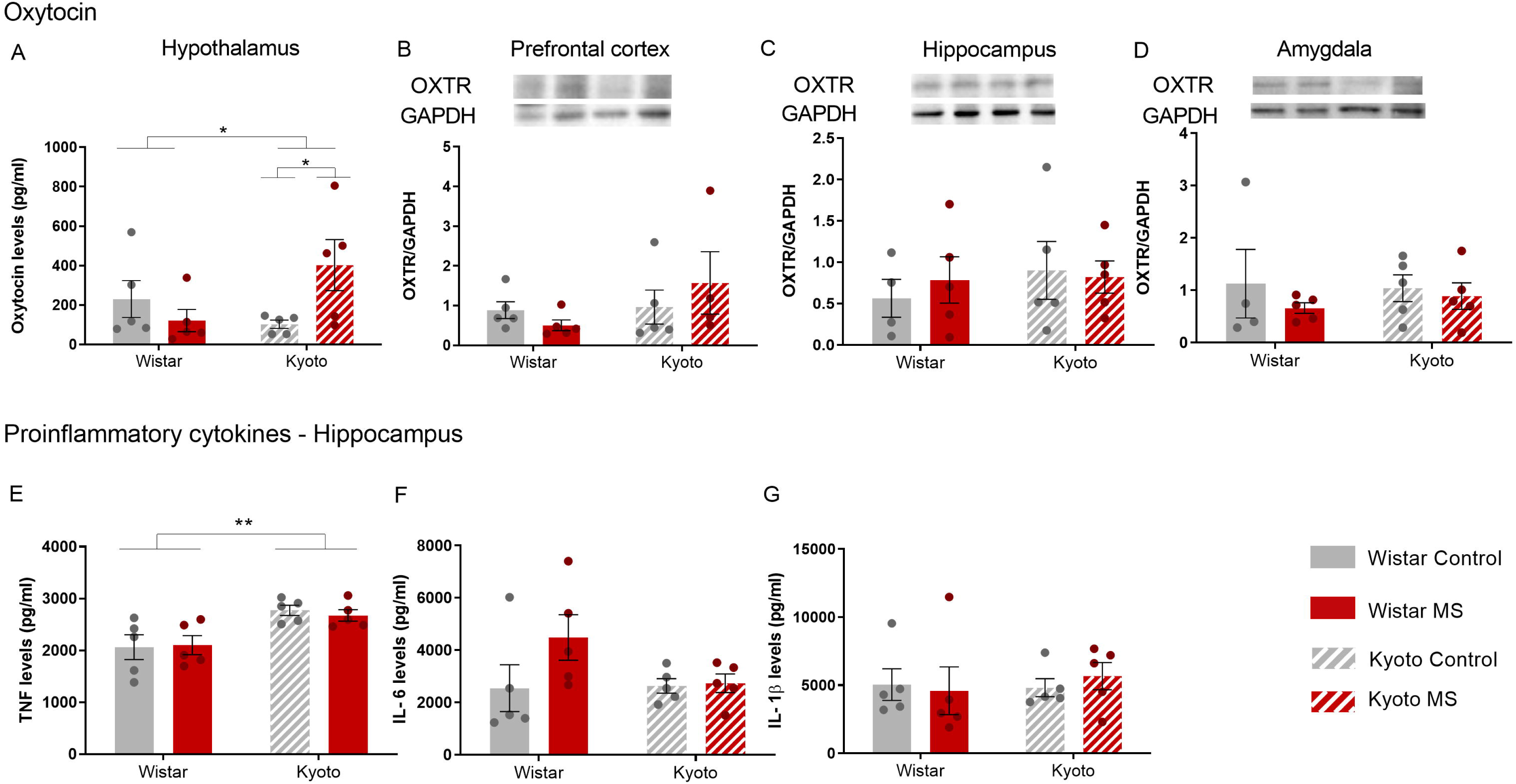
MS disturbed mothers’ neurobehaviour. After weaning, MS increased Kyoto (A) oxytocin levels, in agreement with the decreased of anxiety observed. (E) Kyoto showed high levels of TNF than Wistar strain. Quantification of (A) oxytocin (pg/ml) levels in hypothalamus by ELISA; OXTR in (B) PFC, (C) hippocampus and (D) amygdala by Western blotting; (E) TNF (pg/ml), (F) IL-6 (pg/ml) and (G) IL-1β (pg/ml) levels in hippocampus by ELISA. Data were analysed by two-way ANOVA (2 x 2 factorial design) with Bonferroni comparisons. (A), (B), (C), (F), (G) data were log-transformed. *n* = 5, except for (B) MS Kyoto, and (C) (D) for control Wistar: *n* = 4. Results are expressed as mean ±SEM. \**p* < 0.05, \*\**p* < 0.01, \*\*\**p* < 0.001. MS = Maternal Separation.

#### Kyoto dams present higher TNF levels in the hippocampus than Wistar dams

As proinflammatory cytokines can influence anxiety- and depressive-like behaviours ^18, 19^, we analysed the expression of TNF, IL-6 and IL-1β in the hippocampus ^25,26^and observed that Kyoto dams presented higher expression of TNF in comparison with Wistar dams (*Wald χ*^2^ = 18.3, *p* < 0.001), but no influence of MS **(Fig. 3E)**. Concerning IL-6 **(Fig. 3F)** and IL-1β **(Fig. 3G)** no significant differences between groups were observed.

### Offspring evaluation in response to maternal separation (MS)

#### MS worsened the spatial learning and the reference memory in Kyoto adolescents

To obtain a better understanding of the impact of MS in the adolescent brain, we evaluated the offspring cognitive ability using the Morris Water Maze (MWM). We found that Kyoto adolescents displayed reduced spatial learning in comparison with Wistar (*t*_(348)_ = 7.94, *p* < 0.001). This was aggravated under MS in Kyoto rats, resulting in e an even worse performance in this task (as seen in the comparison with Kyoto controls (strain x MS: *t* = 2.53, *p* < 0.05; and strain x days: *t* = 6.34, *p* < 0.001) **(Fig. 4A)**. During the probe day **(Fig. 4B)**, we also observed that Kyoto performed poorly when compared with Wistar and that the MS worsened both strains regarding the mean distance to the target (strain, *F*_(1,84)_ = 70.4, *p ≤* 0.001, and MS, *F*_(1,84)_ = 9.00, *p* < 0.01) **(Fig. 4C)**. The MS Kyoto and MS Wistar adolescents presented higher mean distance to target in comparison to their respective control groups (*p* < 0.05) **(Fig. 4C)**. Concerning other measurements obtained in the probe day, we observed that the Kyoto strain displayed a worse performance when compared with Wistar. Kyoto adolescents spent less time in the target quadrant (*Wald χ*^2^ = 12.7, *p* < 0.001) **(Fig. 4D)**, presented a higher latency to enter in the target quadrant (*Wald χ*^2^ = 13.9, *p* < 0.001) **(Fig. 4E),** and reduced the number of target crossings (*Wald χ^2^* = 15.4, *p* < 0.001) **(fig. 4F)**. No MS effects were observed in these parameters. Regarding the total distance travelled, we observed that the Kyoto strain travelled less than Wistar and the MS Kyoto adolescents travelled even less (strain, *F*_(1,84)_ = 106, *p <*0.001, and MS: *F*_(1,84)_ = 5.03, *p* < 0.05) **(Fig. 4G)**. For working memory evaluation, we found again differences between strains (*t*_(346)_ = 17.7, *p* < 0.001) but without MS-induced changes **(Fig. 4H)**. Altogether, we observed that Kyoto adolescents presented reduced spatial learning, reference and working memory in comparison with Wistar. MS worsened Kyoto spatial learning and reference memory performance.

**Figure 4.**
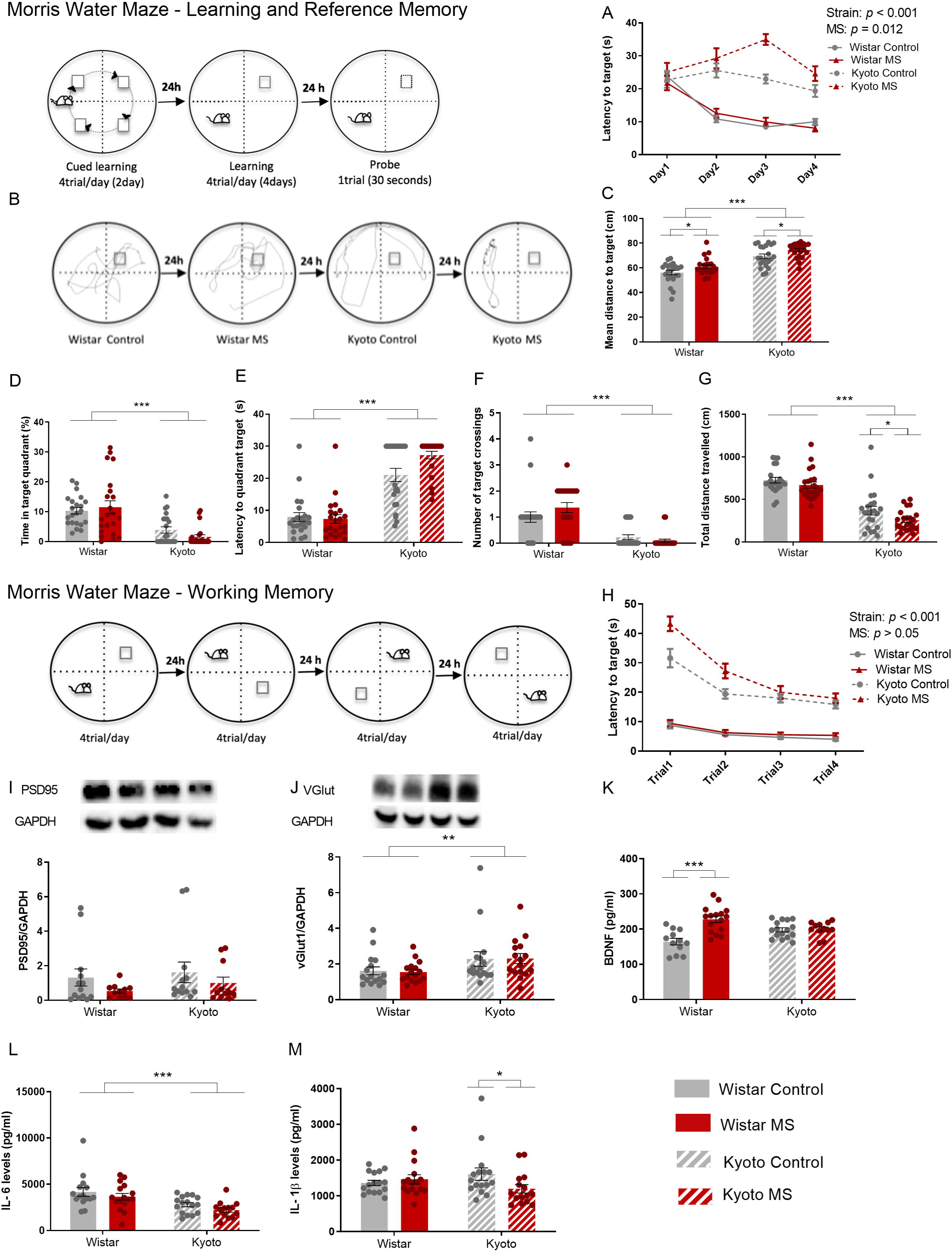
MS worsened Kyoto adolescent offspring cognitive performance. MS increased Kyoto (A) latency to reach the platform, showing learning impairment. Regarding probe day (C), (D), (E), (F) and (G) MS slightly affected both strains. (H) MS did not impact working memory. (J) VGlut expression was higher in Kyoto strain. (K) MS led to an increase of BDNF level on Wistar, which may be related to the increase of affiliative behaviour received by their mothers. (M) MS altered IL1-β levels on Kyoto that can be associated with cognitive impairments presented. Quantification of (I) PSD95 and (J) VGlut in hippocampus by Western blotting; (K) BDNF (pg/ml), (L) IL-6 (pg/ml) and (M) IL-1β (pg/ml) levels in hippocampus by ELISA. (A), (H) data were analysed by linear regression. (C), (G), (I), (J), (K), (L) data were analysed by two-way ANOVA (2 x 2 factorial design) with Bonferroni comparations. (D), (E), (F) data were analysed by logistic regression. (I), (J), (M) data were log-transformed. (A), (C), (D), (E), (F) and (G) *n* = 22. (I), (J), (K), (L), (M) *n* = 16, except for (I) *n* = 13 for control Wistar and Kyoto, *n* = 11 for MS Wistar and Kyoto; (K) *n* = 12 for control Wistar and MS Kyoto, (L) *n* = 15 for control Wistar and *n* = 14 for MS Kyoto, (M) *n* = 15 for MS Kyoto. Results are expressed as mean ±SEM. \**p* < 0.05, \*\**p* < 0.01, \*\*\**p* < 0.001. MS = Maternal Separation; PND = Postnatal day.

#### Kyoto adolescents presented higher levels of vGlut1 expression

To better understand the learning and memory deficits that were observed in the behavioural tests, we analysed the expression of two different glutamatergic synaptic markers in the hippocampus. The postsynaptic density protein 95 (PSD95), is a well-characterized and abundant scaffold protein present in the glutamatergic postsynaptic terminal ^27^, for which no differences in hippocampal expression were observed **(Fig. 4I)**. The vesicular glutamate transporter 1 (vGlut1), is a presynaptic transporter protein that mediates glutamate uptake into synaptic vesicles, the expression level of these transporters determines the vesicle load in glutamate, thereby shaping the efficacy of glutamatergic neurotransmission ^28^. We found that Kyoto adolescents presented a higher expression of vGlut1 (*F*_(1,60)_ = 7.91 *p <* 0.01) **(Fig. 4J)**. Of note, either increased or decreased levels of vGlut1 are associated with impaired cognitive performance ^29, 30^.

#### MS increased BDNF expression in Wistar but not in Kyoto adolescents

As Brain Derived Neurotrophic Factor (BDNF) is a crucial mediator of neuronal plasticity, and a key factor for synapse formation and maintenance during development, as well as for memory processes ^31^, we decided to evaluate BDNF expression in the hippocampus of Wistar and Kyoto rats exposed or not to MS. We observed that BDNF expression was significantly impacted by MS (*F*_(1,52)_ = 17.2, *p < 0*.001). We observed also and also a strain x MS interaction (*F*_(1,52)_ = 14.9, *p <* 0.001), which reflected an MS-induced increase in BDNF expression in Wistar adolescents when compared with control Wistar (*p <* 0.001) **(Fig. 4K)**. This increase in hippocampal BDNF following MS was also observed in previous studies, which proposed to explain it using the ‘mismatch hypothesis’, i.e., that early aversive experiences may trigger adaptive processes, allowing better adaptation to later aversive events ^32^. However, it is also possible that the increased BDNF levels in MS Wistar adolescents may result from the improved maternal care reported in **Fig. 1**.

#### MS decreased IL-1β expression in Kyoto adolescents

Because proinflammatory cytokines can influence learning, memory and shape the neuronal circuit ^33, 34^ and also because altered BDNF levels may associate with altered inflammatory markers ^35^, we decided to evaluate the IL-6 and IL-1β expression in the hippocampus of Wistar and Kyoto rats exposed or not to MS. We observed that Kyoto adolescents presented lower levels of IL-6 than Wistar adolescents (*F*_(1,57)_ = 17.1, *p* < 0.001) **(Fig. 4L)**. For IL-1β, we found a strain x MS interaction (*F*_(1,59)_ = 4.50, *p <* 0.05), corresponding to MS-decreased expression of IL-1β levels in Kyoto adolescents (*p* < 0.05). No MS effects were observed for Wistar adolescents **(Fig. 4M)**. Taken together, these results seem to indicate that a worse cognitive performance was associated with lower levels of pro-inflammatory markers. Previous studies have in fact determined that reduced levels of IL-1β, are associated with hippocampal-dependent learning deficits in the Morris water maze ^36^.

## Discussion

This study investigated the impact of MS on maternal behaviour of dams with different vulnerabilities to depressive-like behaviour and explored its consequences on adolescent cognitive performance. Considering the high comorbidity between motheŕs depression and low maternal care ^1^, we predicted that MS would reduce maternal behaviour in Kyoto mothers. Surprisingly, our results showed an increase in maternal behaviour in both strains in response to MS with different maternal behaviour characteristics after the mother-pups reunion, demonstrating that each strain had different strategies for handling this stressful situation (MS): depressive-like mothers (Kyoto), spent more time in contact (simple physical contact) with pups, whereas non-depressive mothers (Wistar) spent more time in affiliative behaviours (licking/grooming and nursing the pups). During simple physical contact, mothers were lying over the pups (skin contact), but without any oriented behaviour toward the offspring. This skin contact between mother and offspring is important for the acquisition of the pup huddling preference for a maternally-associated odour ^37, 38^. On the other hand, licking/grooming and nursing behaviours are the most common pup-oriented maternal behaviours ^39^. These affiliative maternal behaviours are common in mammals and are crucial for mother-pup attachment ^40^

Our results showed that the maternal depressive-like behaviour traits played an important role with regards to how each dam adjusted its maternal behaviour in response to parental stress (MS). MS Kyoto dams exhibited a more passive strategy towards their offspring, suggesting a lower quality of maternal care in comparison with MS Wistar dams. However, the differences observed in maternal care after MS did not compromise the pups weight gain (a basic measure of growth efficiency). These results further support a previous study that revealed no differences between Wistar and Kyoto offspring’s physical development after MS ^22^. Champagne ^41^ also reported that offspring of low licking mothers do not differ in weaning weight and survival rate. These results suggest that even though the Kyoto maternal behaviour appears to lack quality, these mothers exhibit awareness towards their pups and do not compromise the offspring’s physical health.

Variations in maternal care are associated with differences in oxytocin and OXTR expression in the brain ^42^. Although we did not find alterations in OXTR, surprisingly, we showed for the first time that when a depressive-like dam is exposed to parenting stress (MS), the oxytocin expression by the hypothalamus is augmented. In accordance, previous findings reported that skin contact between mother and pups, but not maternal licking/grooming, was positively related to pups’ hypothalamic oxytocin ^37, 38^, which, in turn, can also be happening in mothers and help explain our observations.

Taking ours and Kojima and colleagues ^43^ results together, it seems that the release of oxytocin by both, mother and offspring, evokes social synchrony. This social synchrony in an adverse situation, here represented by MS, may contribute to better-synchronized interactions, promoting social cohesion, as a defensive behaviour toward potential threats. Thus, the increased levels of oxytocin through maternal contact, in depressive-like mothers in response to MS, may promote cohesion between mother-offspring.

Moreover, the increase in central oxytocin observed in depressive mothers could enhance sensitivity for social salience ^44^, which may increase the sensibility to negative social cues related to the offspring absence. This intensification of the attention to the negative social cues would elicit a more protective behaviour from depressive mothers, serving as defence against a potentially dangerous intruder. The mechanism by which oxytocin affects social behaviour depends on contextual and individual factors ^45^, which seems to influence the sensitivity and interpretation of the emotional meaning or relevance of a situation.

MS depressive mothers, in combination with the higher levels of oxytocin in the hypothalamus, showed a decrease in anxiety-like behaviour in the OF and an increase in the number of arm entries in the EPM. These combined results reflect disinhibition of exploratory behaviour as a consequence of lower anxiety in MS Kyoto dams. This outcome supports studies reporting an oxytocin anxiolytic effect ^46,47^and studies that report an increase in central oxytocin in response to conditioned fear and restrain ^48, 49^, both stressful situations as the MS paradigm used in the present work. In agreement, Light and colleagues ^50^ suggested that oxytocin released during stressful situations serves to alleviate physiological stress. The decreased anxiety-like behaviour observed in depressive-like mothers can be a result of increased oxytocin central levels.

Wistar mothers increased licking/grooming and nursing behaviours due to MS with no impact in the oxytocinergic system. These results suggest that Wistar dams increased pro-social behaviour to counteract the MS effects, exhibiting a more positive emotional regulation in coping the parenting stress situation. Regarding the anxiety-like behaviour, evaluated by EPM and OF tests, Wistar dams did not reveal changes in response to MS. Interestingly, Wistar dams presented increased exploratory behaviours, here represented by rearing and head dipping, that were increased in response to MS.

Of note, maternal behaviour was also evaluated by the PRT. In this test, MS did not affect the retrieval behaviour. This result contrasted with maternal behaviour focal observations. This seeming inconsistency is also observed in other studies. Many of them suggest that PRT should not be used to measure maternal behaviour ^4, 51^ since the PRT is a better predictor of mothers’ activity in the OF. In fact, our combined OF/PRT data support this statement showing that the dams OF activity, like PRT, was not affected by MS.

Moreover, the lack of increased oxytocin expression in the hypothalamus in the MS Wistar mothers corroborates the idea that Wistar mothers were more emotionally regulated, given that higher oxytocin levels in the hypothalamus are part of the physiological response to social stress ^50^ Interestingly, attenuation of the anxiety response by oxytocin occurs in individuals with poor coping, but not in individuals with adequate coping ^52, 53^, what was in accordance with our observations for the Kyoto and Wistar strains.

In the absence of other comorbidities, depressive disorders are related to an increase in central and peripheral proinflammatory cytokines, and mostly with TNF ^19^. In agreement, we observed that TNF levels were higher in depressive-like Kyoto dams in comparison with Wistar. However, there is also a large number of studies reporting that oxytocin reduces TNF production, inducing an anti-inflammatory state ^54^. However, the mechanisms involved are elusive and in vitro results seem to indicate that oxytocin does not directly regulate TNF ^54^. As such, it is possible that the higher TNF levels observed are constitutive in Kyoto dams, and do not seem to respond to increased oxytocin levels.

Maternal care critically affects offspring’s brain maturation, behaviour, cognitive and emotional in several mammalian species ^55^. We observed that the depressive-like adolescent offspring (Kyoto) had significant learning and memory deficits. Also, MS induced a slight impairment in reference memory in both strains, however, the impact of MS was higher in the depressive-like offspring. These results suggest that MS effects on cognition depend on the interaction between genetic (depressive trait) and environmental factors (maternal care quality and early life stress, here represented by MS). Thus, non-depressive-like adolescent offspring raised by higher licking and grooming mothers show only a slight MS impact on cognitive performance.

We also evaluated the expression of synaptic proteins in the hippocampus (major role in learning and memory) and observed a higher vGlut1 expression in Kyoto adolescents, which is in accordance with the memory deficits shown in the behavioural test. Our data highlight vGlut1 importance in memory as observed in a previous aging rat model study in which the increase in vGlut1 expression was related to deficits in spatial memory ^30^

Wistar adolescents exposed to MS had an increased BDNF expression. As BDNF is a crucial factor for learning and memory, the BDNF augment in response to MS in Wistar could have a protective role against MS, blocking the MS-induced learning deficits. Besides, high levels of licking and grooming (maternal behaviour quality) have been associated with higher levels of BDNF in offspring later ^56^, which explains not only the increased BDNF expression in Wistar in response to MS and its protective role regarding cognitive deficits, but also the worse cognition observed in Kyoto adolescents in response to MS.

Pro-inflammatory cytokines are critical factors for memory and cognition, and at either increased or decreased levels they can be detrimental to brain function. IL-6 is an important cytokine for spatial learning and reference memory formation ^33, 57^. In agreement, Kyoto adolescents displayed reduced spatial learning and reference memory in comparison with Wistar, which is in line with the reduced IL-6 expression observed in that strain. However, further studies will be necessary to determine the nature of such decreased IL-6 levels.

Adequate levels of IL-1β are necessary for learning and memory processes ^58^. We found IL-1β expression decreased in the hippocampus of MS Kyoto adolescents, which performed poorly in the MWM. In accordance, mice lacking the IL-1 receptor present hippocampus-dependent learning deficits reflected in poorer performance also in the MWM ^36^, showing that endogenous IL-1β is necessary for learning and memory. Likewise, the administration of an IL-1 receptor antagonist (IL-1ra) also causes spatial memory impairment ^59^.

Considering our initial predictions, our findings were somewhat unexpected since the results presented here cannot be explained by the notion that higher oxytocin levels increase pro-social behaviours. Our study evidenced that beyond pro-social behaviours, oxytocin is involved in stress and mood disorder regulation (**Fig. 5**). We showed for the first time that when a depressive mother is exposed to parenting stress, the release of endogenous hypothalamic oxytocin is triggered and leads to an altered perception of the environment (as insecure). In addition, in combination with the studies of Kojima and colleagues ^37, 38, 43^, our study suggests that oxytocin not only increases sensitivity to environmental cues, but it may also amplify the social stress transmission contributing to a bidirectional intensification of social synchrony. This social synchrony in an adverse situation may subsequently contribute to better interactions, enabling the refinement of social communication and consequently promoting social cohesion as a defensive behaviour toward potential threats. Finally, our results also demonstrate that different maternal stress-coping styles affect the offspring behaviour, providing important clues about how early life stress increases the risk for poor cognitive performances and how maternal care is a possible target for intervention.

**Figure 5.**
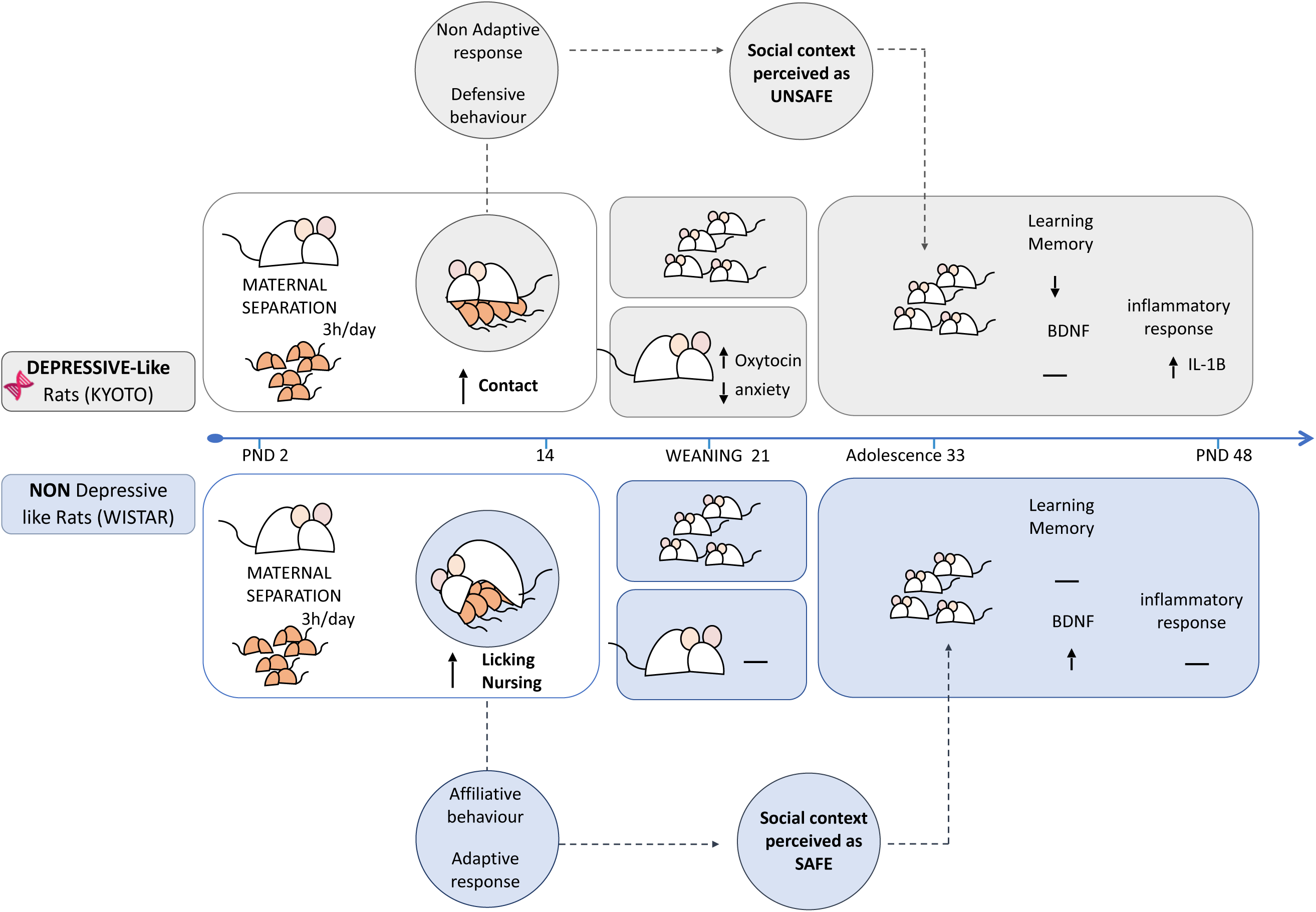
Graphical summary of main findings. Our results showed that mothers with different depression-like vulnerabilities facing parental stress (induced by MS) adopted different strategies. Although both strains increased the maternal behaviour in response to MS, non-depressive-like mothers (Wistar) exhibited a higher quality maternal behaviour (affiliative), showing a more adaptive response and more pro-social behaviour (more active strategy). On the other hand, depressive-like mothers (Kyoto) displayed a more passive/defensive strategy (non-adaptive response) in response to MS. The higher levels of oxytocin observed in Kyoto mothers that experienced MS seem to be part of the stress-response mechanism, which amplifies the negative perception of the environment (as insecure), leading to a more defensive behaviour by keeping the pups very close to them and away from outsiders (out-group anti-social behaviour). This increase in oxytocin expression seems to decrease the anxiety-like state in Kyoto dams in response to MS. Furthermore, the quality of maternal behaviour observed in non-depressive-like mothers (Wistar) after MS seems to protect the cognitive performance of adolescent rats from the negative effects of MS, leading to an increase of BDNF expression in the offspring hippocampus. On the other hand, depressive-like adolescents showed lower resilience to MS effects, exhibiting worst performance in the cognitive tests and alteration in the IL-1β levels.

## Methods

### Animals

This study used Wistar and Kyoto rats originally obtained from Charles River (Barcelona/Spain). Rats were kept under standard conditions (21±1°C; 60±5% humidity; 12h light/12h dark cycle - light off at 12h00). Food and water were available *ad libitum.* Mating was done by introducing a male into a cage with two females (nulliparous) at the beginning of the dark cycle. On gestation day 16, pregnant females were individually housed. Nest material was provided to each dam and no bedding changes were performed in the last days of pregnancy. All experiments involving animals were approved by the Portuguese regulatory agency Direcção Geral de Alimentação e Veterinária (DGAV) and the animal ethics committee of IBMC-i3S. The animal facility and the people directly involved in animal experimentation were also certified by DGAV. All animal experiments considered the Russell and Burch 3R’s principle and followed the European guidelines for animal welfare (2010/63/ EU Directive). All time measurements were recorded in seconds and distances in centimetres.

### Maternal separation (MS) protocol

The delivery day was designated as postnatal day (PND) 0. Pups were sexed and each litter was adjusted to 8-10 pups in PND 1. Dams from each strain were randomly assigned to control (*n* = 8/strain) or to the MS group (*n* = 7/strain). Animals from the control group were undisturbed except for cage change once a week (animal facility reared). The MS group dams were separated from their offspring at 9:00 am, 180min per day, from PND 2-14. During the separation period, litters and mothers were placed in a different cage and in different rooms to avoid ultrasonic communication. Hypothermia during MS was prevented by placing the pup cages on a heating pad. In PND 21, rats were weaned and co-housed (2-3 per cage) and the following groups were established (*n* = 11 per group): (a) Wistar male control group; (b) Wistar female control group; (c) Kyoto male control group; (d) Kyoto female control group; (e) Wistar male MS group; (f) Wistar female MS group; (g) Kyoto male MS group; and (h) Kyoto female MS group. Each group contained rats from all litters to avoid possible confounding litter effects.

### Behavioural test

To evaluate the impact of MS in mother’s behaviour, maternal behaviour (at PND 2, 6, 10 and 14) and PRT (PND 4) were evaluated soon after pup reunion. In addition, to study if MS exacerbates the anxiety- and depressive-like behaviour profile of dams, the mothers were subjected to the EPM (PND 7) during MS and to the OF (PND 21) and FST (PND 22) after weaning. At PND 23, brain areas were collected to examine the oxytocinergic system and proinflammatory cytokines. Then, when offspring reached the adolescent period (PND 28-50), the effects of MS on learning and memory were evaluated by MWM (PND 37). At PND 48, in order to study memory and learning targets, the offspring hippocampi were collected **(Fig.6)**.

**Figure 6.**
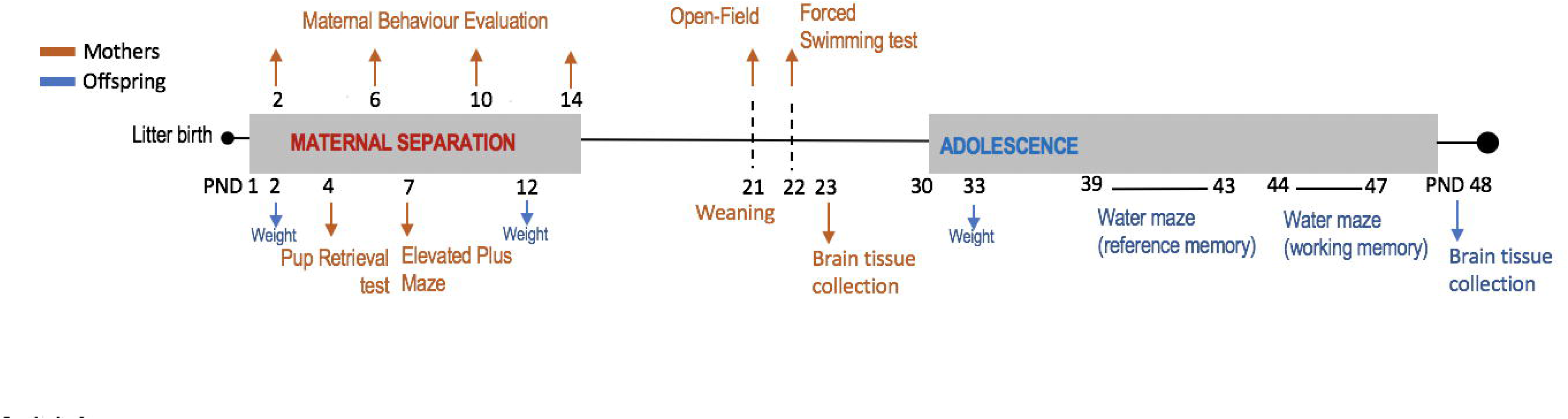
Schematic timeline summarizing the experimental procedures. PND = Postnatal day.

### Mother’s behaviour evaluation

#### Maternal behaviour

To evaluate the effects of MS on the quantity and quality of maternal care, maternal behaviours toward offspring were scored immediately after pup reunion, on PND 2, 6, 10 and 14, for 10 min. Animals were not disturbed during behavioural observations. Maternal behaviours were divided into the following categories: (a) total maternal behaviour (sum of affiliative and non-affiliative contact behaviour); (b) affiliative behaviour - time spent grooming/licking and nursing; and (c) non-affiliative behaviour - time dams spent in contact with the offspring but not in grooming/licking and nursing. Behaviour was analysed using the *focal-animal* sampling method with the software Observer XT version10 (Noldus information technology, Netherlands).

#### Pup Retrieval Test (PRT)

The PRT was performed at PND 4 to complement the maternal behaviour analysis. After MS, pups were returned to the original cage at the opposite side of the nest (far from the nest), followed by the return of the mother. The control group cages had mothers removed for less than 3 min and returned, during this time pups were moved to the opposite nest site. The observation ended when mothers retrieved all pups, or the 10 min observation limit was achieved ^60^. The following behaviours were quantified using the software Observer XT version10 (Noldus information technology, Netherlands): (a) latency to first contact, (b) latency to first pup retrieval, and (c) full litter retrieval duration.

#### Elevated Plus Maze (EPM)

Mother anxiety-like behaviour was measured during the MS period (PND 7). The apparatus consisted of four connected arms (44 cm x 14 cm) in a plus-shaped structure and elevated 72 cm above the floor. Two arms were an open platform (no sidewalls) and two were enclosed within 22cm-high sidewalls. Arms were positioned so opposing arms were of the same type. Each mother was placed in the central platform (the intersection of the four arms - 14×14 cm area) facing an open arm and left undisturbed for 5 min. Testing was conducted in the dark phase of the light/dark cycle. The apparatus was cleaned between animals with neutral, odour free soap. The following behaviours were measured using the software Observer XT version 10 (Noldus information technology, Netherlands): (a) open arms time; (b) number of open arm entries; (c) centre area time; (d) immobility time; (e) total number of arm entries; (f) number of rearing; and (g) number of head dipping.

#### Open-Field (OF)

On weaning day (PND 21), mothers were tested in the OF to measure anxiety-like symptoms. The apparatus consisted of an empty rectangular arena (80 x 60 cm). Mothers were placed in the corner of the apparatus and left to spontaneously explore the arena for 10 min. The following behaviours were automatically recorded by the Smart Video Tracking Software (Panlab) version 3.0: (a) time in centre area; (b) number of centre entries; (c) latency to the centre zone; (d) total distance travelled; and (e) number of rearing.

#### Forced Swimming Test (FST)

Mother depressed-like behaviour was assessed with the FST at PND 22 according to the protocol described by Aguggia and colleagues ^60^. Dams were placed in a transparent container filled with water (25±2°C) for 10 min (pre-test session to reduce acute stress) and tested 24 hours later for 5 min in the same container. The following behaviours were quantified by the Software Observer XT version10 (Noldus information technology, Netherlands): (a) time spent climbing and (b) time spent in immobility, defined as the time spent either immobile or making righting movements to stay afloat.

### Adolescent offspring behaviour evaluation

#### Pup’s weight

Pup’s weight was measured at PND 2, 12 and 33.

#### Morris Water Maze (MWM)

The MWM test was used to evaluate the impact of MS on the cognitive performance of the adolescent offspring. The procedure used was adapted from the one previously described by Vorhees and Williams ^61^. The MWM apparatus consisted of a circular pool (111 cm diameter) filled with water (24°C), divided into 4 imaginary equal-sized quadrants created by two lines (N-S and W-E) perpendicular to each other and with the intersection at the centre of the pool. The quadrants were named SW, SE, NW, NE. A square platform, made of transparent acrylic, was placed in the centre of one of the quadrants. To provide external references, multiple extra-maze visual cues of different shapes were hung on the wall of the experimental room. Indirect lighting was provided.

In preparation for the memory task tests, cued learning was done at PND 37 and 38. The extra-maze visual cues were made not visible (exclusive to cued learning days) and the platform was slightly submerged in the swimming pool with an object on top (visible escape platform). Four trials were performed during which both, start quadrant and platform locations were changed. If a rat did not locate the platform before 60 s elapsed, it was guided to the platform and was allowed to stay there for 15 s.

#### Spatial learning and reference memory task

During the next four days after cued learning (PND 39-42), rats were subjected to the spatial learning task (four trials per day). Extra maze cues were placed in the room and the platform was submerged (near the surface). The platform was placed in a fixed location (same for all spatial learning trials) and the rats were gently released into the tank at one of the four starting positions and allowed to locate the hidden escape platform for a maximum of 60 s. When rats failed to find the platform, the experimenter guided the animal to it. Rats were allowed to stay on the platform for 15 s. Latency to reach the platform was recorded. On probe day (PND 43), the reference memory was evaluated (1 trial). The platform was removed, and rats were released in the quadrant opposite the platform previous location. Rats were allowed to swim freely during 30 s. The following measures were quantified by the Smart Video Tracking Software (Panlab) version 3.0: (a) mean distance to target; (b) total distance travelled; (c) percentage of total time in the target quadrant; (d) the latency to the target quadrant; and (e) number of target crossings.

#### Spatial working memory task

At PND 44-47 the MWM was used to study working memory (four trials per day). In this protocol (adapted from Tata and colleagues^62^), the platform and rat starting positions were the same in each day but changed every day. Maze cues were kept as in the spatial learning task. Thus, there was no platform location learning from one day to the next day, but latency to reach the platform is expected to decrease within the day, from trial one to four. As in the spatial learning task, rats had 60 s to reach the escape platform; if animals failed, the experimenter guided them to the platform, where they were allowed to stay for 15 s. Latency to reach the platform was recorded.

### Mothers and adolescent offspring biomarkers evaluation

#### Protein extract preparation

Mother’s hypothalamus, amygdala, prefrontal cortex (PFC) and hippocampus were homogenized in a buffer containing 50 mM Tris-Base, 10 mM EDTA, 150 mM NaCl, 0.1% Tween 20 and protease inhibitors ^63^. The offspring hippocampi were homogenized in a buffer containing 50 mM Tris-Base, 4 mM EDTA, 150 mM NaCl, 0.1% Tween 20 and protease inhibitors. Samples were sonicated and centrifuged for 15 min at 2271×g and posteriorly, the supernatant was centrifuged for 5 min at 13500×g. These centrifugations allowed the removal of large aggregates ^63^. The concentration of protein was estimated using Pierce BCA Protein Assay kit (Thermo Scientific), following the manufacturer’s instructions.

#### Enzyme-Linked Immunosorbent Assay (ELISA)

ELISA was performed using the Oxytocin ELISA kit (ADI-900-153A; Enzo Life Sciences) to measure the oxytocin levels on mother’s hypothalamus. A total of 40 µg/well of hypothalamus protein were used and samples were analysed in duplicates. The BDNF of offspring hippocampus was measured using the ChemiKineTM BDNF Sandwich ELISA kit (CYT306; Chemikine, Millipore). A total of 80 µg/well of hippocampus protein was used and samples were analysed in duplicates. The quantification of IL-1β, IL-6 and TNF in the mother’s hippocampus and the quantification of IL-1β and IL-6 in the offspring’s hippocampus were performed by ELISA (Rat IL-1β Mini ABTS ELISA Development Kit: 900-M91; Rat IL-6 Mini ABTS ELISA Development Kit: 900-M86; Rat TNF-α Mini ABST ELISA Development Kit: 900-M73; ABTS ELISA Buffer Kit: 900-K00 - PeproTech). A total of 80 µg/well of hippocampus protein was used. All the assays were performed by the manufacturer’s instructions and using a microplate reader (Tecan - Sunrise).

#### Western Blot

The expression of OXTR on mother’s PFC, hippocampus (using 100 µg sample) and amygdala (using 70 µg sample) was analysed by Western Blot. Western Blot was also used to detect the expression of the PSD95 and vGlut1 on offspring’s hippocampus (using 30 µg sample). Samples were denatured at 95 °C for 5 min and were loaded into a 10% SDS-page gel. Proteins were transferred onto a nitrocellulose membrane using Trans-Blot® Turbo™ RTA Midi Nitrocellulose Transfer Kit (Bio-Rad) in the Trans-Blot® Turbo™ Transfer System (Bio-Rad). Membranes were blocked for 1 hour and incubated with primary antibody overnight, at 4 °C (oxytocin receptor (Thermo Fisher Scientific): 1:400; PSD95 (Thermo Fisher Scientific): 1:2000; vGlut1 (Synaptic Systems): 1:1000; GAPDH (HyTest): 1:100000), and with secondary antibody for 1 hour, at room temperature (HRP conjugated anti-rabbit 1:10000 and HRP conjugated anti-mouse 1:10000 (both from Jackson ImmunoResearch). Membranes were developed using SuperSignal™ West Pico PLUS Chemiluminescent Substrate (Thermo Fisher Scientific) in the ChemiDoc (ChemiDoc™ MP System, Bio-Rad). Results were quantified using Fiji – ImageJ software.

### Statistical analyses

Normality was verified by the Shapiro-Wilk or Kolmogorov-Smirnov test. When measures did not validate normality assumption but verified the homogeneity of variances, results were accepted assuming the robustness of the ANOVA test. The assumption of homogeneity of variance was verified by Levene’s test. Regarding mother data, a two-way ANOVA (2×2 factorial design) was used, with strain (Wistar, Kyoto), and MS treatment (control, MS) as independent factors. Concerning adolescent data, a three-way ANOVA (2×2×2 factorial design) was initially conducted, adding sex as an independent factor. However, since sex was found not significant (*p* > 0.05), it was excluded, and a two-way ANOVA was conducted. A Linear Mixed Model (LMM) was used to verify if litter effect (as a random variable) was significant for the offspring measures, but significance was not found (except for pup weight). Bonferroni test was used to perform multiple comparisons. When some measures did not validate the assumption of homogeneity, all measures were transformed in accordance (percentages were arcsine transformed; measures with zero values were square-root transformed and the other cases log-transformed). In the PRT - latency to first pup return, EPM-number of open arms entries, EPM – number of total arm entries, OF – latency normality assumption was violated even with transformation. EPM - open arm time, EPM - centre area time, EPM - number of head dipping, and TNF levels violated the assumption of homogeneity of variances even with data transformation. A Generalized Linear Model with a robust estimator was conducted in these cases with the transformed values due to normality validation. Litter size effect in PRT results was also considered as a random effect in an LMM, but since there was no significant statistical effect of litter size (*p* = 0.588), the LMM was dropped and results were based on the ANOVA. In MWM, spatial learning and working memory were first analysed by a Repeated Measures ANOVA with *days* (spatial learning at day 1, 2, 3, and 4) or *trails* (working memory at trial 1, 2, 3, and 4) as within-subject factor, and strain (Wistar, Kyoto) and MS treatment (control, MS) as between-subject factor. However, since homogeneity of variances by Levene’s test was violated, a linear regression with log-transformation was conducted. In addition, in MWM, time in target quadrant, number of target crossings and latency to target quadrant data could not be analysed with an ANOVA due to assumption violation; thus, a binary variable was created (0 - not going to; 1 - going to) and a logistic regression test was used to determine the effects of strain and MS treatment. IBM SPSS Statistics 23 software was used to analyse all data. Alpha level for statistical significance was set at 0.05.

## Acknowledgements

This work was financed by FEDER - Fundo Europeu de Desenvolvimento Regional funds through the COMPETE 2020 - Operational Programme for Competitiveness and Internationalisation (POCI), Portugal 2020, and by Portuguese funds through FCT - Fundação para a Ciência e a Tecnologia/Ministério da Ciência, Tecnologia e Ensino Superior in the framework of the project POCI-01-0145-FEDER-032231 (PTDC/SAU- TOX/32231/2017) and by FCT and Orçamento do Estado in the framework of the project EXPL-AMAGALHÃES - IF/00753/2014/CP1241/CT0005. RLA was supported by an FCT grant (PD/BD/114266/2016). CCP holds an employment contract financed by national funds through FCT - Fundação para a Ciência e a Tecnologia, I.P., in the context of the program-contract described in paragraphs 4, 5 and 6 of art. 23 of Law no. 57/2016, of August 29, as amended by Law no. 57/2017 of July 19. AM was supported by FCT (IF/00753/2014).

## Author Contributions

RLA, AM conceived the study and edited the paper. RLA, AM and CCP conceived the experimental approach. RLA, CCP, CJA, AM conducted the experiments and RLA, PO, IML analysed the results. RLA, CCP, IML, FB, TS, AM co-wrote the manuscript. All authors critically discussed the results and reviewed the final version of the manuscript.

## References

1. Righetti-Veltema, M., Conne-Perreard, E., Bousquet, A. & Manzano, J. Postpartum depression and mother-infant relationship at 3 months old. J Affect Disord 70, 291–306 (2002).

2. Vedam, S. et al. The Mothers on Respect (MOR) index: measuring quality, safety, and human rights in childbirth. SSM Popul Health 3, 201–210, doi:10.1016/j.ssmph.2017.01.005 (2017).

3. Fleming, A. S., O’Day, D. H. & Kraemer, G. W. Neurobiology of mother–infant interactions: experience and central nervous system plasticity across development and generations. Neuroscience & Biobehavioral Reviews 23, 673–685, doi:https://doi.org/10.1016/S0149-7634(99)00011-1 (1999).

4. Alves, R. L., Portugal, C. C., Summavielle, T., Barbosa, F. & Magalhaes, A. Maternal separation effects on mother rodents’ behaviour: A systematic review. Neurosci Biobehav Rev, doi:10.1016/j.neubiorev.2019.09.008 (2019).

5. Insel, T. R. & Young, L. J. The neurobiology of attachment. Nature Reviews Neuroscience 2, 129–136, doi:10.1038/35053579 (2001).

6. Meyer-Lindenberg, A., Domes, G., Kirsch, P. & Heinrichs, M. Oxytocin and vasopressin in the human brain: social neuropeptides for translational medicine. Nat Rev Neurosci 12, 524–538, doi:10.1038/nrn3044 (2011).

7. Pedersen, C. A. & Boccia, M. L. Oxytocin antagonism alters rat dams’ oral grooming and upright posturing over pups. Physiology & behavior 80, 233–241, doi:10.1016/j.physbeh.2003.07.011 (2003).

8. Ferris, C. F. in Progress in Brain Research Vol. 170 (eds Inga D. Neumann & Rainer Landgraf) 305–320 (Elsevier, 2008).

9. Amini-Khoei, H. et al. Oxytocin mitigated the depressive-like behaviors of maternal separation stress through modulating mitochondrial function and neuroinflammation. Progress in Neuro-Psychopharmacology and Biological Psychiatry 76, 169–178, doi:https://doi.org/10.1016/j.pnpbp.2017.02.022 (2017).

10. Newport, D. J., Stowe, Z. N. & Nemeroff, C. B. Parental depression: animal models of an adverse life event. The American journal of psychiatry 159, 1265–1283, doi:10.1176/appi.ajp.159.8.1265 (2002).

11. Kaidbey, J. H. et al. Early Life Maternal Separation and Maternal Behaviour Modulate Acoustic Characteristics of Rat Pup Ultrasonic Vocalizations. Scientific Reports 9, 19012, doi:10.1038/s41598-019-54800-z (2019).

12. Champagne, F., Diorio, J., Sharma, S. & Meaney, M. J. Naturally occurring variations in maternal behavior in the rat are associated with differences in estrogen-inducible central oxytocin receptors. Proceedings of the National Academy of Sciences 98, 12736, doi:10.1073/pnas.221224598 (2001).

13. Holubová, A., Lukášková, I., Tomášová, N., Šuhajdová, M. & Šlamberová, R. Early Postnatal Stress Impairs Cognitive Functions of Male Rats Persisting Until Adulthood. Frontiers in Behavioral Neuroscience 12, doi:10.3389/fnbeh.2018.00176 (2018).

14. Huot, R. L., Plotsky, P. M., Lenox, R. H. & McNamara, R. K. Neonatal maternal separation reduces hippocampal mossy fiber density in adult Long Evans rats. Brain research 950, 52–63, doi:10.1016/s0006-8993(02)02985-2 (2002).

15. Aisa, Tordera, R., Lasheras, B., Del Rio, J. & Ramirez, M. J. Effects of maternal separation on hypothalamic-pituitary-adrenal responses, cognition and vulnerability to stress in adult female rats. Neuroscience 154, 1218–1226, doi:10.1016/j.neuroscience.2008.05.011 (2008).

16. Calcia, M. A. et al. Stress and neuroinflammation: a systematic review of the effects of stress on microglia and the implications for mental illness. Psychopharmacology 233, 1637–1650, doi:10.1007/s00213-016-4218-9 (2016).

17. Kumar, A. Editorial: Neuroinflammation and Cognition. Front Aging Neurosci 10, 413–413, doi:10.3389/fnagi.2018.00413 (2018).

18. Hurley, L. L. & Tizabi, Y. Neuroinflammation, neurodegeneration, and depression. Neurotoxicity research 23, 131–144, doi:10.1007/s12640-012-9348-1 (2013).

19. Dowlati, Y. et al. A meta-analysis of cytokines in major depression. Biological psychiatry 67, 446–457, doi:10.1016/j.biopsych.2009.09.033 (2010).

20. Himmerich, H. et al. Depression, comorbidities and the TNF-alpha system. European psychiatry : the journal of the Association of European Psychiatrists 23, 421–429, doi:10.1016/j.eurpsy.2008.03.013 (2008).

21. Voorhees, J. L. et al. Prolonged restraint stress increases IL-6, reduces IL- 10, and causes persistent depressive-like behavior that is reversed by recombinant IL-10. PloS one 8, e58488–e58488, doi:10.1371/journal.pone.0058488 (2013).

22. Rana, S., Pugh, P. C., Jackson, N., Clinton, S. M. & Kerman, I. A. Inborn stress reactivity shapes adult behavioral consequences of early-life maternal separation stress. Neurosci Lett 584, 146–150, doi:10.1016/j.neulet.2014.10.011 (2015).

23. Nam, H., Clinton, S. M., Jackson, N. L. & Kerman, I. A. Learned helplessness and social avoidance in the Wistar-Kyoto rat. Front Behav Neurosci 8, 109, doi:10.3389/fnbeh.2014.00109 (2014).

24. van Zyl, P. J., Dimatelis, J. J. & Russell, V. A. Changes in behavior and ultrasonic vocalizations during antidepressant treatment in the maternally separated Wistar-Kyoto rat model of depression. Metab Brain Dis 29, 495–507, doi:10.1007/s11011-013-9463-6 (2014).

25. Sheline, Y. I., Mittler, B. L. & Mintun, M. A. The hippocampus and depression. European Psychiatry 17, 300–305, doi:https://doi.org/10.1016/S0924-9338(02)00655-7 (2002).

26. Qiao, H. et al. Dendritic Spines in Depression: What We Learned from Animal Models. Neural plasticity 2016, 8056370, doi:10.1155/2016/8056370 (2016).

27. Kim, E. & Sheng, M. PDZ domain proteins of synapses. Nature Reviews Neuroscience 5, 771–781, doi:10.1038/nrn1517 (2004).

28. Wojcik, S. M. et al. An essential role for vesicular glutamate transporter 1 (VGLUT1) in postnatal development and control of quantal size. Proceedings of the National Academy of Sciences of the United States of America 101, 7158, doi:10.1073/pnas.0401764101 (2004).

29. Du, X. et al. Research progress on the role of type I vesicular glutamate transporter (VGLUT1) in nervous system diseases. Cell & bioscience 10, 26, doi:10.1186/s13578-020-00393-4 (2020).

30. Ménard, C. et al. Glutamate presynaptic vesicular transporter and postsynaptic receptor levels correlate with spatial memory status in aging rat models. Neurobiology of aging 36, 1471–1482, doi:10.1016/j.neurobiolaging.2014.11.013 (2015).

31. Yamada, K. & Nabeshima, T. Brain-derived neurotrophic factor/TrkB signaling in memory processes. Journal of pharmacological sciences 91, 267–270, doi:10.1254/jphs.91.267 (2003).

32. Récamier-Carballo, S., Estrada-Camarena, E. & López-Rubalcava, C. Maternal separation induces long-term effects on monoamines and brain-derived neurotrophic factor levels on the frontal cortex, amygdala, and hippocampus: differential effects after a stress challenge. Behavioural pharmacology 28, 545–557, doi:10.1097/fbp.0000000000000324 (2017).

33. Bialuk, I., Taranta, A. & Winnicka, M. M. IL-6 deficiency alters spatial memory in 4- and 24-month-old mice. Neurobiology of learning and memory 155, 21–29, doi:10.1016/j.nlm.2018.06.006 (2018).

34. Huang, Z. B. & Sheng, G. Q. Interleukin-1β with learning and memory. Neuroscience bulletin 26, 455–468, doi:10.1007/s12264-010-6023-5 (2010).

35. Jin, Y., Sun, L. H., Yang, W., Cui, R. J. & Xu, S. B. The Role of BDNF in the Neuroimmune Axis Regulation of Mood Disorders. Frontiers in Neurology 10, 515 (2019).

36. Avital, A. et al. Impaired interleukin-1 signaling is associated with deficits in hippocampal memory processes and neural plasticity. Hippocampus 13, 826–834, doi:10.1002/hipo.10135 (2003).

37. Kojima, S. & Alberts, J. R. Maternal care can rapidly induce an odor-guided huddling preference in rat pups. Developmental psychobiology 51, 95–105, doi:https://doi.org/10.1002/dev.20349 (2009).

38. Kojima, S., Stewart, R. A., Demas, G. E. & Alberts, J. R. Maternal contact differentially modulates central and peripheral oxytocin in rat pups during a brief regime of mother-pup interaction that induces a filial huddling preference. Journal of neuroendocrinology 24, 831–840, doi:10.1111/j.1365-2826.2012.02280.x (2012).

39. Orso, R. et al. How Early Life Stress Impact Maternal Care: A Systematic Review of Rodent Studies. Frontiers in Behavioral Neuroscience 13, doi:10.3389/fnbeh.2019.00197 (2019).

40. Keverne & Curley. Vasopressin, oxytocin and social behaviour Current Opinion in Neurobiology 14, 777–783, https://doi.org/10.1016/j.conb.2004.10.006 (2004).

41. Champagne, F. A., Francis, D. D., Mar, A. & Meaney, M. J. Variations in maternal care in the rat as a mediating influence for the effects of environment on development. Physiology & behavior 79, 359–371, doi:10.1016/s0031-9384(03)00149-5 (2003).

42. Rilling, J. K. & Young, L. J. The biology of mammalian parenting and its effect on offspring social development. Science 345, 771–776, doi:10.1126/science.1252723 (2014).

43. Kojima, S. & Alberts, J. R. Warmth from skin-to-skin contact with mother is essential for the acquisition of filial huddling preference in preweanling rats. Developmental psychobiology 53, 813–827, doi:10.1002/dev.20565 (2011).

44. Olff, M. et al. The role of oxytocin in social bonding, stress regulation and mental health: an update on the moderating effects of context and interindividual differences. Psychoneuroendocrinology 38, 1883–1894, doi:10.1016/j.psyneuen.2013.06.019 (2013).

45. Bartz, J. A., Zaki, J., Bolger, N. & Ochsner, K. N. Social effects of oxytocin in humans: context and person matter. Trends Cogn Sci 15, 301–309, doi:10.1016/j.tics.2011.05.002 (2011).

46. Bale, T. L., Davis, A. M., Auger, A. P., Dorsa, D. M. & McCarthy, M. M. CNS region-specific oxytocin receptor expression: importance in regulation of anxiety and sex behavior. The Journal of neuroscience : the official journal of the Society for Neuroscience 21, 2546–2552, doi:10.1523/jneurosci.21-07-02546.2001 (2001).

47. Ring, R. H. et al. Anxiolytic-like activity of oxytocin in male mice: behavioral and autonomic evidence, therapeutic implications. Psychopharmacology 185, 218–225, doi:10.1007/s00213-005-0293-z (2006).

48. Onaka, T. Neural pathways controlling central and peripheral oxytocin release during stress. Journal of neuroendocrinology 16, 308–312, doi:10.1111/j.0953-8194.2004.01186.x (2004).

49. Neumann, I. D., Krömer, S. A., Toschi, N. & Ebner, K. Brain oxytocin inhibits the (re)activity of the hypothalamo-pituitary-adrenal axis in male rats: involvement of hypothalamic and limbic brain regions. Regulatory peptides 96, 31–38, doi:10.1016/s0167-0115(00)00197-x (2000).

50. Light, K. C. et al. Deficits in plasma oxytocin responses and increased negative affect, stress, and blood pressure in mothers with cocaine exposure during pregnancy. Addict Behav 29, 1541–1564, doi:10.1016/j.addbeh.2004.02.062 (2004).

51. Curley, J. P., Jensen, C. L., Franks, B. & Champagne, F. A. Variation in maternal and anxiety-like behavior associated with discrete patterns of oxytocin and vasopressin 1a receptor density in the lateral septum. Hormones and behavior 61, 454–461, doi:10.1016/j.yhbeh.2012.01.013 (2012).

52. Cardoso, C., Linnen, A. M., Joober, R. & Ellenbogen, M. A. Coping style moderates the effect of intranasal oxytocin on the mood response to interpersonal stress. Experimental and clinical psychopharmacology 20, 84–91, doi:10.1037/a0025763 (2012).

53. Quirin, M., Kuhl, J. & Düsing, R. Oxytocin buffers cortisol responses to stress in individuals with impaired emotion regulation abilities. Psychoneuroendocrinology 36, 898–904, doi:10.1016/j.psyneuen.2010.12.005 (2011).

54. Panaro, M. A., Benameur, T. & Porro, C. Hypothalamic Neuropeptide Brain Protection: Focus on Oxytocin. Journal of Clinical Medicine 9, doi:10.3390/jcm9051534 (2020).

55. Bath, K., Manzano-Nieves, G. & Goodwill, H. Early life stress accelerates behavioral and neural maturation of the hippocampus in male mice. Hormones and behavior 82, 64–71, doi:10.1016/j.yhbeh.2016.04.010 (2016).

56. Liu, Diorio, Day, Francis & Meaney. Maternal care, hippocampal synaptogenesis and cognitive development in rats. Nature neuroscience 3, 799–806, doi:10.1038/77702 (2000).

57. Baier, P. C., May, U., Scheller, J., Rose-John, S. & Schiffelholz, T. Impaired hippocampus-dependent and -independent learning in IL-6 deficient mice. Behav Brain Res 200, 192–196 (2009).

58. Brennan, F. X., Beck, K. D. & Servatius, R. J. Low doses of interleukin-1β improve the leverpress avoidance performance of Sprague–Dawley rats. Neurobiology of learning and memory 80, 168–171, doi:https://doi.org/10.1016/S1074-7427(03)00060-1 (2003).

59. Yirmiya, R., Winocur, G. & Goshen, I. Brain Interleukin-1 Is Involved in Spatial Memory and Passive Avoidance Conditioning. Neurobiology of learning and memory 78, 379–389, doi:https://doi.org/10.1006/nlme.2002.4072 (2002).

60. Aguggia, J. P., Suarez, M. M. & Rivarola, M. A. Early maternal separation: neurobehavioral consequences in mother rats. Behav Brain Res 248, 25–31, doi:10.1016/j.bbr.2013.03.040 (2013).

61. Vorhees & Williams. Morris water maze: procedures for assessing spatial and related forms of learning and memory. Nature protocols 1, 848–858, doi:10.1038/nprot.2006.116 (2006).

62. Tata. Maternal separation as a model of early stress: Effects on aspects of emotional behavior and neuroendocrine function. Hellenic Journal of Psychology 9, 84–101 (2012).

63. Clarke, M. & Pearl, C. A. Alterations in the estrogen environment of the testis contribute to declining sperm production in aging rats. Systems biology in reproductive medicine 60, 89–97, doi:10.3109/19396368.2014.885995 (2014).

